# Phirefly: hierarchical quantum haplotype phasing at the chromosome scale

**DOI:** 10.64898/2026.11.24.768929

**Authors:** Yifei Wang, Megan K Le, Heng Li

## Abstract

**Motivation:** Read-based phasing links alleles on the same homologous haplotype. It plays a critical role in long-read variant calling and variant analysis. Many phasing algorithms have been developed. This includes QHap, the only quantum-inspired algorithm for read-based phasing. QHap does not run on quantum hardware due to the scale of the practical problems.

**Results:** We developed Phirefly, the first practical quantum algorithm for read-based phasing with the key phasing step executed on quantum hardware. Phirefly uses a CPU-based algorithm to identify small disjoint groups of SNPs linked by strong, consistent read support. These groups are called phaselets and are easy to phase. To phase across phaselets, which is harder due to weak or conflicting read support, Phirefly locks the relative SNP phases within each phaselet and applies a quantum algorithm, effectively treating each phaselet as one heterozygous variant. Phirefly runs on physical quantum computers from D-Wave and IBM, as well as with a quantum-inspired classical solver using CPU or GPU. It achieves broadly comparable accuracy to QHap and mainstream phasing algorithms on HG002 datasets.

**Availability and implementation:** https://github.com/wangyf1125/Phirefly

## 1. Introduction

Haplotype phasing is the process of determining which alleles occur on the same chromosome copy. It supports the interpretation of genetic variation and the reconstruction of haplotype-resolved genomes. Long reads link heterozygous variants within individual DNA molecules, providing direct evidence for their relative phase. However, sequencing and alignment errors can introduce conflicting phase relationships among reads, which read-based phasing methods must reconcile to reconstruct haplotypes.

Read-based phasing is often formulated as the weighted minimum error correction (wMEC) problem, which is an NP-hard optimization problem. WhatsHap solves this formulation through dynamic programming, whereas HapCUT2 optimizes a likelihood model using graph cuts (Patterson et al., 2015; Edge et al., 2017). They are the two most widely used algorithms. Many other methods exist as well: HapCompass uses spanning trees and conflicting cycles, HAHap assembles local haplotypes hierarchically, HapCHAT merges reads, and SpecHap uses spectral graph analysis (Aguiar and Istrail, 2012; Lin et al., 2018; Beretta et al., 2018; Yu et al., 2021). We recently published longcallR that jointly infers read haplotags and SNP phases for long-read RNA sequencing (Huang et al., 2026). This will form the basis of our Phirefly algorithm.

Read-based phasing can also be formulated as an Ising model (Tourdot et al., 2021), which underlies a large number of problems amendable by quantum algorithms (Farhi et al., 2014). QHap takes such a model and is the first quantum-inspired algorithm for read-based phasing (Zhang et al., 2026). However, it would not work on quantum hardware due to limited qubit availability in physical quantum processors (Preskill, 2018). There are no quantum phasing algorithms.

In this article, we introduce Phirefly for long-read haplotype phasing on quantum computers (Figure 1). It is the first quantum algorithm that works for chromosome-scale phasing. We applied Phirefly to real HG002 datasets with the MindQuantum (Xu et al., 2024) simulator using GPU, and also on IBM gate-based processors and a D-Wave quantum annealer. We evaluated its accuracy in comparison to QHap and the two mainstream phasing algorithms, WhatsHap and HapCUT2.

**Figure 1.**
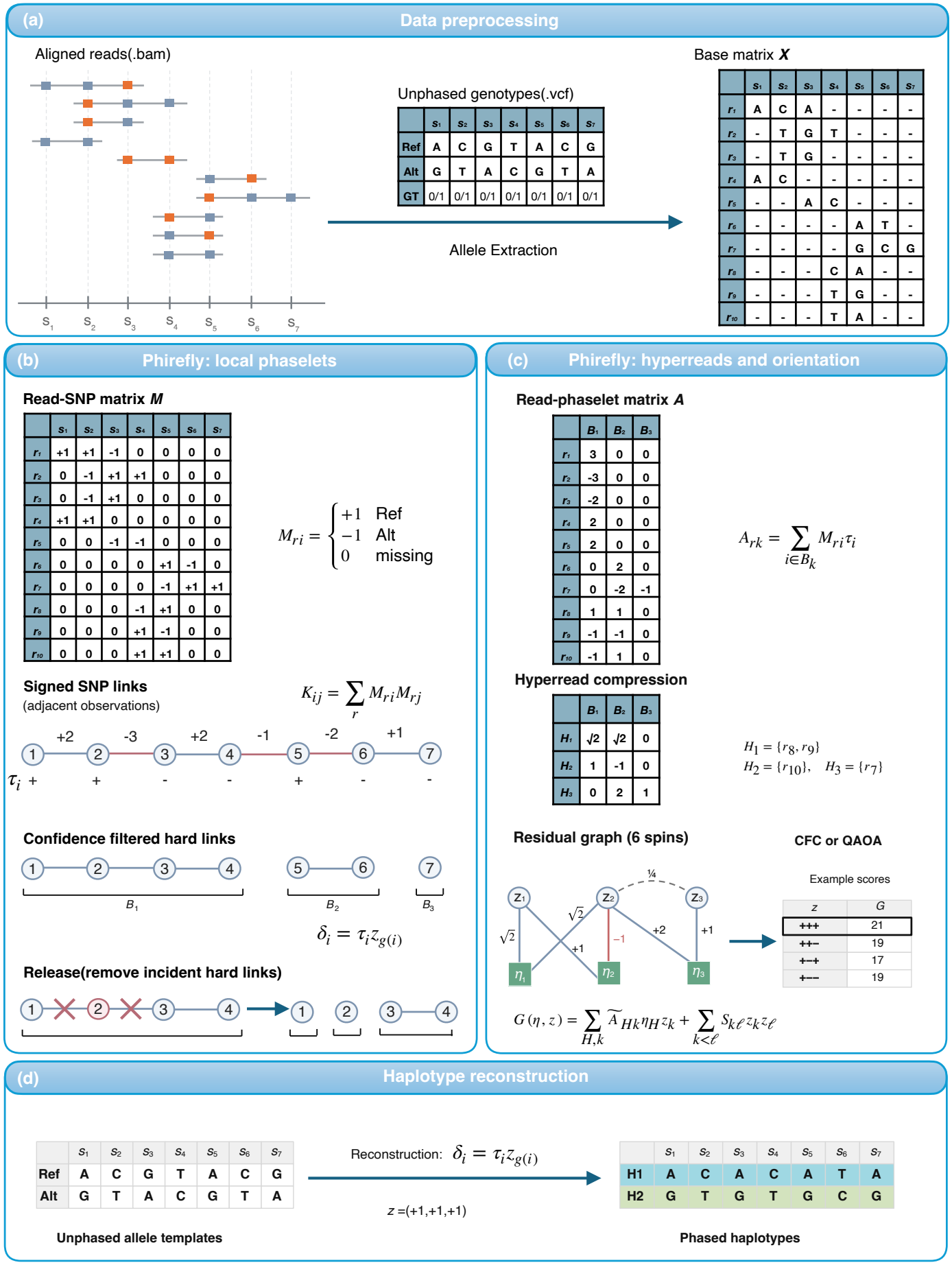
Haplotype phasing with Phirefly. (a) A read contributes one allele observation at each heterozygous SNP it covers. (b) A maximum-spanning forest assigns initial SNP signs. Strong links group SNPs into phaselets. Phirefly removes links at sites where the reads favor the opposite sign. (c) A read that supports only one phaselet has the same best score in either orientation. We group reads that span the same phaselets with the same relative signs into a hyperread. For each connected component, the solver samples phaselet orientations *z* and hyperread labels *η*. We keep the sampled pair (*η, z*) with the lowest energy, *E* = −*G*(*η, z*). (d) The selected orientation of phaselet *g*(*i*) gives the phase of SNP *i* as *δ*_*i*_ = *τ*_*i*_*z*_*g*(*i*)_.

## 2. Materials and methods

Phirefly takes aligned long reads in the BAM format and unphased heterozygous SNP genotypes in the VCF format as input and writes a phased VCF file (Figure 1). Phirefly first constructs a graph with SNPs as vertices and read supports as edge weights. It then finds a maximum-spanning forest (MSF) on the graph and selects edges with strong, consistent read support. The selected edges after this procedure group SNPs into connected components, which we call *phaselets*. To reduce search space, Phirefly combines reads that link the same phaselets and support the same relative orientations into *hyperreads*. Phirefly then builds an Ising model with two types of variables: one for each phaselet orientation and the other for each hyperread haplotag. To find a solution to the Ising model, we use chaotic feedback control (CFC), a classical quantum-inspired algorithm, on a GPU, the quantum approximate optimization algorithm (QAOA) on an IBM quantum processor, or quantum annealing on a D-Wave processor. For each connected component of the Ising graph, we use the phaselet orientations from the sample that best agrees with the reads to assign SNP phases.

### 2.1. Read–SNP model for haplotype phasing

We consider diploid phasing at fixed biallelic heterozygous SNPs. For each observed read–SNP pair (*r, i*) ∈ O, let *b*_*ri*_ = +1 indicate the reference allele and *b*_*ri*_ = −1 the alternate allele. Let *σ*_*r*_ ∈ {−1, +1} label the haplotype of read *r* and let *δ*_*i*_ ∈ {−1, +1} denote the allele on haplotype +1 at SNP *i*: +1 for the reference allele and −1 for the alternate allele. An observation agrees with its assigned haplotype when *b*_*ri*_ = *σ*_*r*_*δ*_*i*_. Maximizing the agreement score

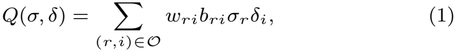

where *w*_*ri*_ ≥ 0 is the observation weight, is equivalent to solving the wMEC problem. Weight *w*_*ri*_ can be calculated from base and mapping quality, though by default, Phirefly sets *w*_*ri*_ = 1 and requires a base quality of at least 13 and a mapping quality of at least 20 because base and mapping quality estimates may not be precise.

Given real data, there are hundreds of thousands of reads and SNPs on a long human chromosome. It is impractical to solve this equation on physical quantum computers.

### 2.2. Constructing phaselets

We construct a signed SNP graph G_*s*_ to initialize relative SNP phases. For each read, we order the observed SNPs on it by genomic position and add an edge, or *link*, between adjacent SNPs on the read. Note that a SNP may be skipped on the read if the read base at the SNP position has base quality below 13, a deletion or a non-SNP base. As a result, G_*s*_ may contain a link between two SNPs not adjacent on the genome, which happens for about a third of links in our MHC test region.

Let ℛ_*ij*_ be the set of reads that contribute a link between SNPs *i* and *j*. With unit observation weights, the signed support for this link is

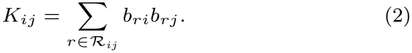

Positive support favors the same SNP signs, whereas negative support favors opposite signs.

We exclude zero-support links and construct a maximum-spanning forest using Kruskal’s algorithm (Kruskal, 1956). We process links in decreasing order of |*K*_*ij*_ | and retain a link only if it connects different components.

With MSF, we can propagate SNP signs along links in the trees. Let *τ*_*i*_ ∈ {−1, +1} be the initial value of *δ*_*i*_ assigned by the forest. We assign +1 to each tree root and isolated SNPs. When we know *τ*_*i*_, we set *τ*_*j*_ = *τ*_*i*_ · sign(*K*_*ij*_) if the link between *i* and *j* is retained. For a SNP pair with *n*_*ij*_ contributing reads, |*K*_*ij*_ |*/n*_*ij*_ is 1 when all reads agree on the relative phase and 0 when they are evenly split. We introduce a filtering score that is lower for pairs with stronger agreement, more reads and shorter distance (Supplementary Section S2 for the exact definition). We consider a link to be *hard* if at least two reads contribute, the SNPs are at most 1 Mb apart, the filtering score is at most 0.45 and the signed support must agree with the MSF phases (i.e., *K*_*ij*_*τ*_*i*_*τ*_*j*_ *>* 0). Of the SNP links not selected as hard links, we retain those that pass a less stringent filter as *soft links*.

Connected components of the hard links form the initial phaselets. Flipping a whole phaselet cannot correct a relative phase error within it. To obtain high-quality phaselets, we further check each SNP before fixing the phaselets. To check SNP *i*, we first infer the haplotag *u*_*r*\*i*_ of each read *r* covering it, using only the other SNPs on that read. Let *M*_*ri*_ = *w*_*ri*_*b*_*ri*_ for observed pairs and zero otherwise. Using the MSF signs *τ*, we compute

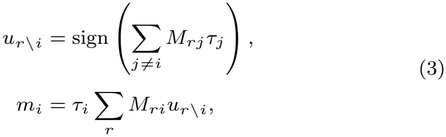

A read contributes nothing if its other SNPs give a tie (*u*_*r*\*i*_ = 0). With unit weights, the margin *m*_*i*_ is the number of reads supporting *τ*_*i*_ minus the number opposing it. If *m*_*i*_ *<* 0, we remove all hard links touching SNP *i*—we call this step *release*. We then rebuild the phaselets from the remaining links, keeping *τ* unchanged. For SNP *i* in the resulting phaselet *g*(*i*), we write

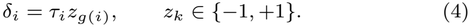

If *z*_*k*_ = +1, the SNPs in phaselet *k* keep their MSF signs; if *z*_*k*_ = −1, they all flip together.

### 2.3. Projecting reads to hyperreads

For each read *r*, we sum its signed observations within phaselet *k*:

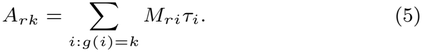

If only one entry *A*_*rk*_ is nonzero, the read contributes |*A*_*rk*_| under either orientation of the phaselet. Therefore we keep this constant and remove the read variable.

For each read that supports at least two phaselets, let *c*_*r*_ be the sign of its first nonzero entry in phaselet order. If this entry is negative, we flip all signs in the row. We then group reads that support the same phaselets with matching signs into a *hyperread H*. The contribution of hyperread *H* to phaselet *k* is

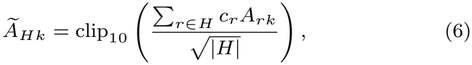

### 2.4. Residual optimization and candidate selection

For two phaselets connected by soft SNP links, we empirically set the weight between them to *S*_*k*ℓ_:

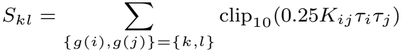

where the sum is carried over soft links between *i* and *j*. The final objective function combines support from hyperreads and soft links:

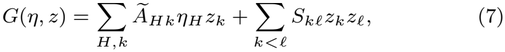

where *η*_*H*_ ∈ {−1, +1} is the label of hyperread *H* and *z*_*k*_ is the orientation of phaselet *k*.

We use CFC on a GPU to sample hyperread labels and phaselet orientations together. By default, we generate one batch of 100 candidates with 1,000 iterations. For each connected component of the residual graph, we keep the sampled assignment with the lowest energy, *E* = −*G*. We use its phaselet orientations to assign SNP phases, *δ*_*i*_ = *τ*_*i*_*z*_*g*(*i*)_, and write one phase set per component to the output VCF.

### 2.5. Hi-C integration

We use Hi-C contacts to infer the relative orientations of phaselets. For each Hi-C alignment that passes filtering, we sum *b*_*ri*_*τ*_*i*_ over the SNPs it covers in each phaselet. We choose the phaselet with the largest absolute sum and record the sign of that sum. Together, the phaselet and sign form an anchor. If all sums are zero, we skip the alignment.

We group alignments by normalized read name and keep the strongest anchor for each phaselet. Each read name contributes at most once to a given phaselet pair. Matching anchor signs favor the same orientation; opposite signs favor opposite orientations. For each pair that passes the support filters, we sum these signed, weighted contributions, multiply by 0.1 and cap the magnitude at 10. The resulting weight *C*_*k*ℓ_ adds a Hi-C term to the long-read score:

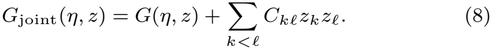

Supplementary Section S6 gives the filters and weights.

We sample the joint model with one CFC batch using the same settings as above. For each component defined by the long-read and Hi-C links, we keep the sample with the lowest energy, *E*_joint_ = −*G*_joint_. Hi-C can change the orientation of a phaselet but not the relative phases of SNPs within it.

### 2.6. Quantum execution

We used the long read graphs from all 22 HG002 autosomes, without Hi-C. We treated each connected component containing at least one edge as a separate problem.

On IBM processors, we used single-layer QAOA (*p* = 1)(Farhi et al., 2014), with one qubit for each phaselet orientation and one for each hyperread label. For couplings *J*_*uv*_ in *G*, the cost Hamiltonian is

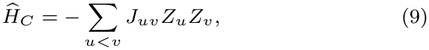

where *Z*_*u*_ is the Pauli-*Z* operator on qubit *u*. For each component, we chose the QAOA angles on a classical computer by minimizing the expected energy and kept them fixed during hardware runs.

IBM sampled 3,903 components with 512 shots per circuit and at most 139 problem qubits. We omitted one 239-variable component on chr19, leaving its 253 observed SNPs unphased. To measure variation across runs, we sampled the same 51 chr22 components three times on each of `ibm_fez`, `ibm_marrakesh` and `ibm_kingston`. The largest chr22 circuit used 67 problem qubits.

On D-Wave `Advantage2_system1`, we sampled all 3,904 components, including the 239 variable component on chr19. We used 512 annealing reads per component and an annealing time of 20 *µ*s per read. We decoded each chain of physical qubits by majority vote.

For the 44 chr22 components with at most 21 variables, we also compared ideal QAOA simulation with uniform sampling of spin assignments. Each sampler generated 512 samples per component for each of 20 seeds.

In all experiments, we kept the sampled assignment (*η, z*) with the lowest energy, *E* = −*G*(*η, z*), in each component. Its phaselet orientations gave the SNP phases, *δ*_*i*_ = *τ*_*i*_*z*_*g*(*i*)_. Supplementary Section S5 gives the angle search, circuit compilation, bit-to-spin mapping and embedding settings. Reported QPU times exclude classical processing and queueing.

### 2.7. Benchmark design

We compared Phirefly, WhatsHap, HapCUT2 and QHap SNP on the 22 HG002 autosomes. All four methods used the same PacBio Revio HiFi reads aligned to GRCh38 at nominal 48-fold coverage and the same 2,180,776 unphased biallelic heterozygous SNP genotypes from the GIAB/NIST v5.0q assembly-based draft benchmark (Genome in a Bottle Consortium, 2023; National Institute of Standards and Technology, 2026).

Each workflow was allocated eight CPU cores and 64 GiB of RAM. WhatsHap and HapCUT2 each ran on a single thread. Phirefly and QHap SNP used up to eight CPU cores for read extraction and one NVIDIA H100 GPU for sampling.

We ran each method three times per chromosome. End-to-end (E2E) time covers the workflow from aligned BAM and unphased VCF input to indexed phased VCF output, including startup, read extraction, graph construction, phasing and export. Core time measures only the solver calls, after their inputs have been built. For GPU methods, Core time includes data transfers to and from the GPU. Final candidate selection and VCF export count only toward E2E. Supplementary Section S3 gives the timing boundaries for each method.

We calculated Hamming error (HE) and switch error (SE) against the reference phases with WhatsHap 2.8. We also report the fraction of input SNPs phased and phase-block N50 and NG50 after splitting overlapping blocks. We average the three runs for each chromosome and report the mean of these 22 values.

## 3. Results

### 3.1. Identifying phaselets

Across the 22 autosomes, Phirefly reduced the number of read and SNP variables by an average factor of 185.7, leaving a mean of 2,177.3 variables per chromosome (Figure 2A,B). CFC sampling accounted for 0.69% of mean end-to-end time (Figure 2D).

**Figure 2.**
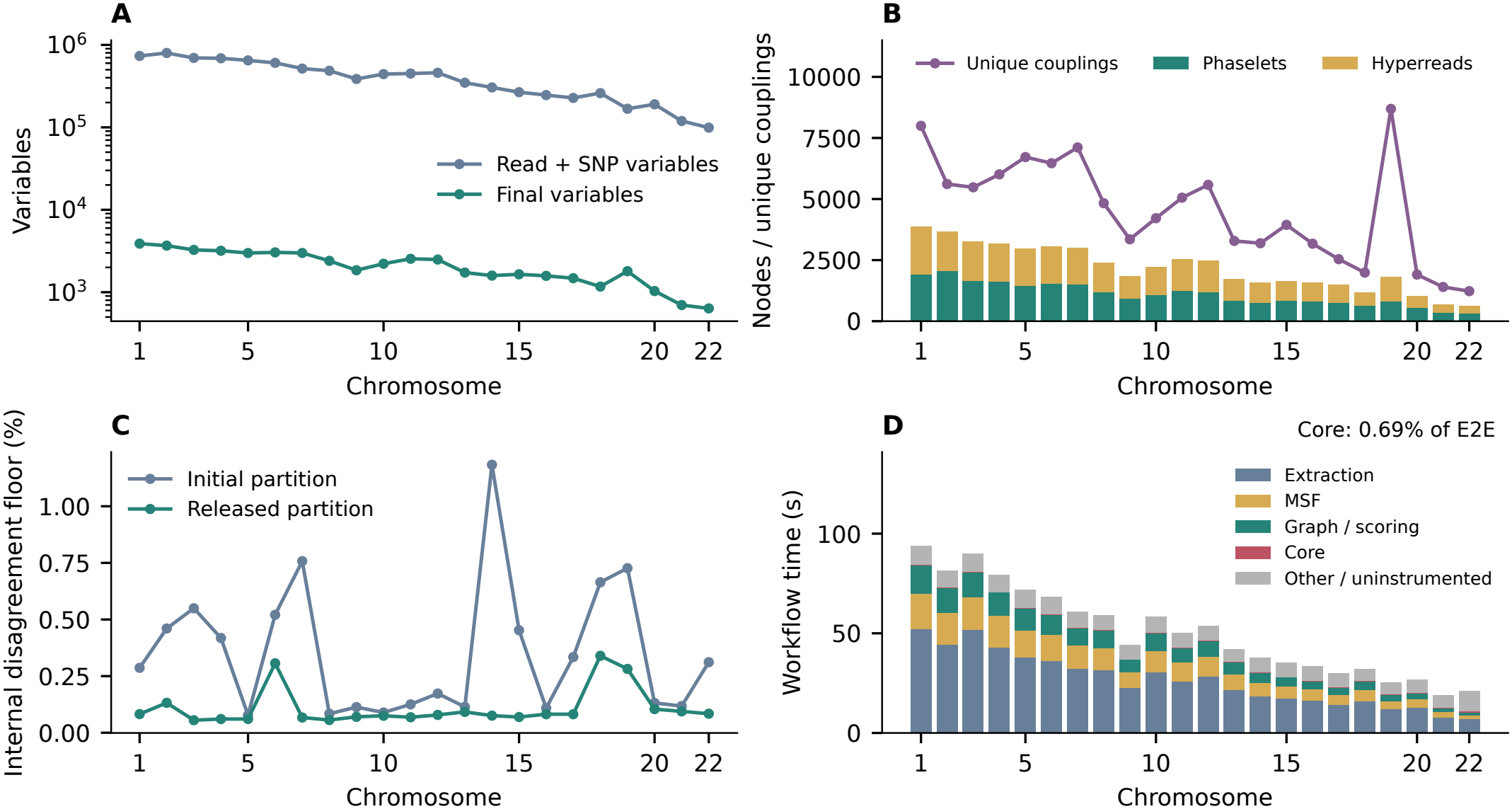
Performance of Phirefly. (A) Variable counts before and after compression on each chromosome. (B) Composition of the compressed graphs. Bars show the numbers of phaselet and hyperread variables; points show the numbers of nonzero couplings, with each pair counted once. (C) Minimum mismatch rate before and after release on each chromosome. Each phaselet was oriented to give the fewest mismatches within each reference block. (D) Mean end-to-end time per chromosome across three runs, broken down by stage.

To count mismatches that a phaselet flip could not correct, we chose the better of the two orientations for each phaselet within each reference block. Among 2,179,974 observed phased SNPs, 7,724 mismatches remained before release and 2,271 after release, a step dropping spurious SNPs (Section 2.2). The mean mismatch rate across chromosomes fell from 0.355% to 0.110% (Figure 2C).

Across all 22 autosomes, we also compared the initial MSF phases with the final Phirefly output using the same observed SNPs and phase sets from the original read graph. Mean HE fell from 5.390% to 0.548%, while mean SE rose from 0.251% to 0.258% (Supplementary Table S3).

### 3.2. Phasing accuracy and complete-workflow time

Phirefly had the shortest end-to-end runtime on the 22 HG002 autosomes, averaging 50.6 s per chromosome, compared with 162.4 s for HapCUT2, 295.4 s for WhatsHap and 312.8 s for QHap SNP

(Table 1; Figure 3). Compared with QHap SNP, Phirefly had lower mean HE (0.548% versus 0.730%) and slightly higher mean SE (0.258% versus 0.248%). Both had a mean PB-N50 of 564 kb. WhatsHap and HapCUT2 had lower HE and SE than Phirefly, with WhatsHap giving the lowest rates overall (0.278% HE and 0.142% SE). Across five CFC seeds, Phirefly’s mean HE ranged from 0.534% to 0.603%, and mean SE from 0.257% to 0.258% (Supplementary Table S2).

**Table 1.**
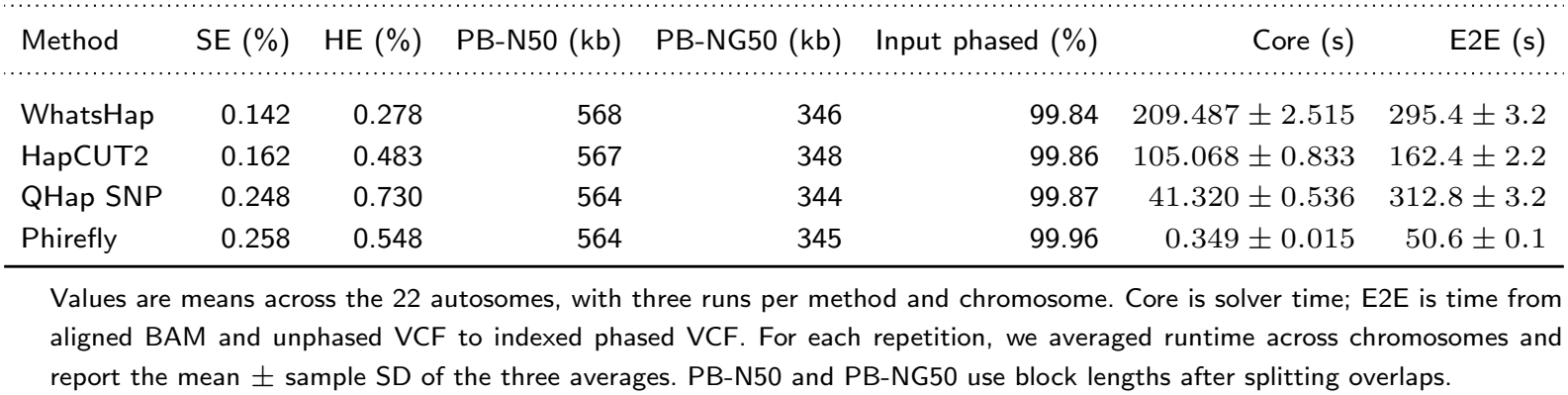
Phasing accuracy and runtime across the 22 HG002 autosomes. Each method ran three times per chromosome.

**Figure 3.**
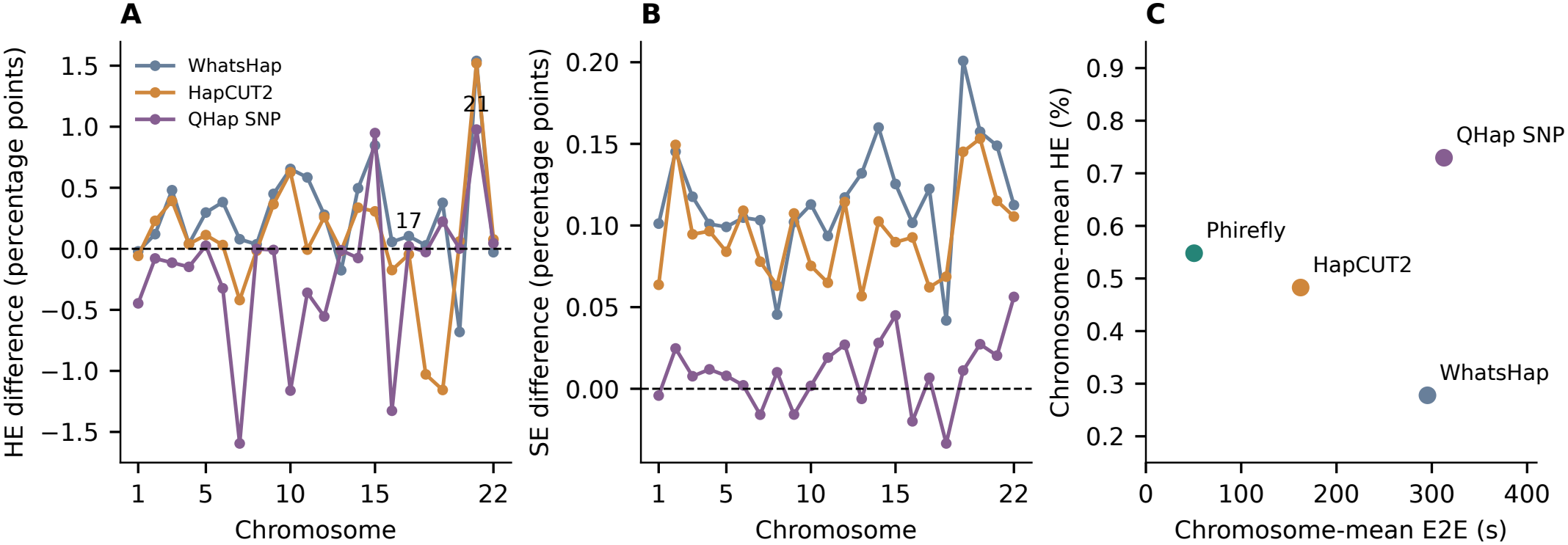
Phasing accuracy and end-to-end runtime. (A,B) Differences in HE and SE, in percentage points, between Phirefly and each comparator on each chromosome. Negative values indicate lower error for Phirefly. (C) Mean HE and end-to-end runtime per chromosome for the four methods.

### 3.3. Hi-C integration

Adding Hi-C to Phirefly increased mean span N50 from 0.564 to 126.0 Mb across the 22 HG002 autosomes. We measured the span of each phase set from its first to its last SNP. Mean HE fell from 0.548% to 0.207%, and mean SE fell from 0.258% to 0.214%.

With the same inputs, HapCUT2 had a mean phase-set span N50 of 115.1 Mb. On their own phase sets, Phirefly had lower HE than HapCUT2 (0.207% versus 0.267%) but higher SE (0.214% versus 0.104%). Because each evaluation block can be oriented independently against the reference, we also compared the methods on the same SNPs and blocks. HapCUT2 then had lower HE (0.120% versus 0.205%) and lower SE on the same adjacent SNP pairs (0.099% versus 0.212%; Supplementary Table S5).

### 3.4. Classical optimization of compressed graphs

We compared seven GPU Ising solvers on the same compressed graphs from chr1, chr20 and chr22 (Figure 4). Each solver used 200 candidates per run and 20 seeds, with schedules of 5,000 or 10,000 iterations. On chr1, LQA came closest to the reference under the 10,000-iteration schedule: its lowest energy per run was 0.972 units above the reference on average. On chr20 and chr22, both BSB and LQA reached the reference in all 20 runs. The reference was the exact minimum of the compressed model for chr22 and the lowest energy found for chr1 and chr20.

**Figure 4.**
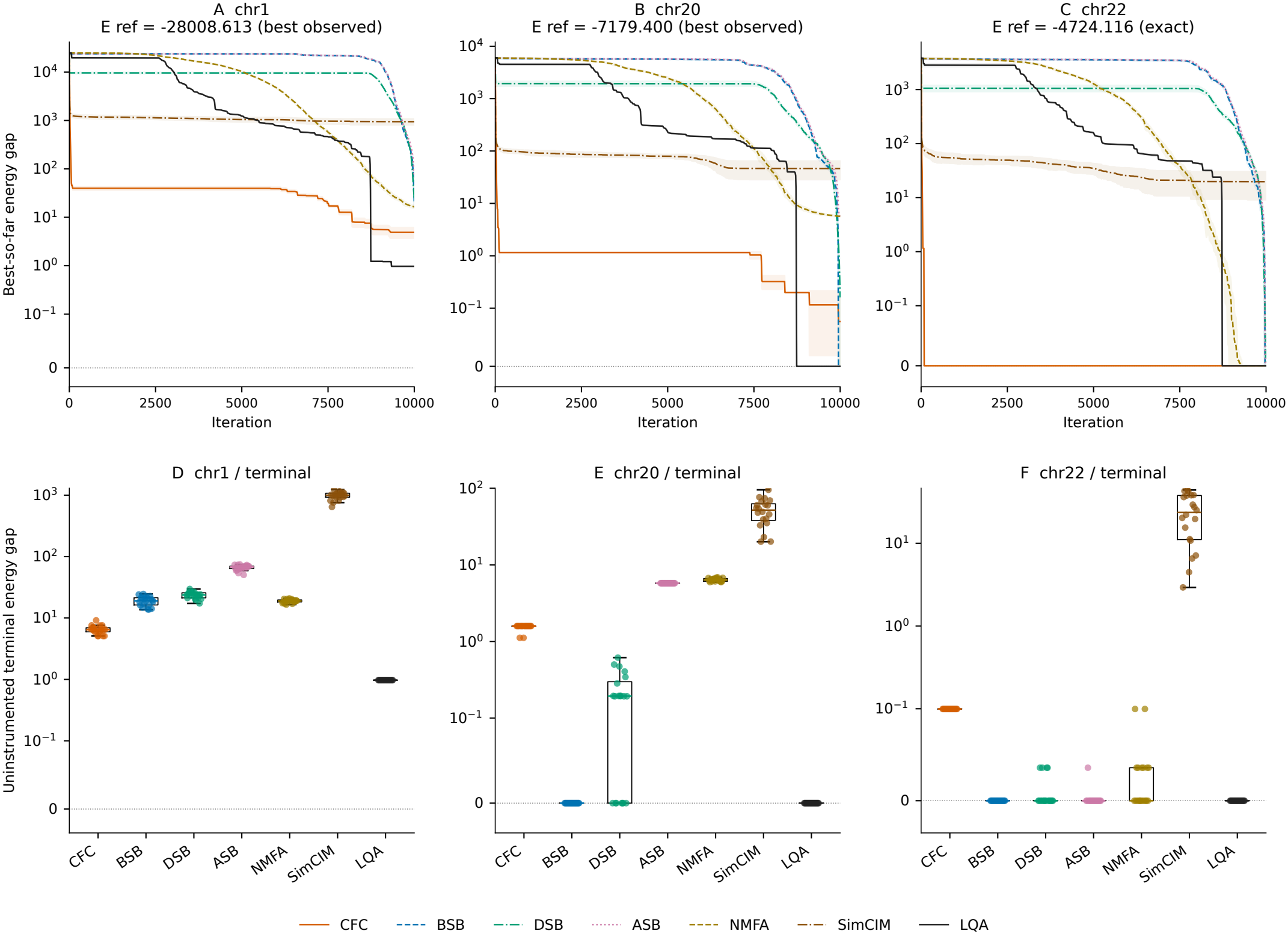
Seven GPU solvers on the compressed chr1, chr20 and chr22 graphs. Energy gaps are measured from the lowest energy found for chr1 and chr20 and the exact minimum for chr22. All panels show the 10,000-iteration schedule with 200 candidates and 20 seeds. (A–C) Lowest energy gap seen so far, recorded every 10 updates and at the endpoint. Lines and shading show the mean and standard deviation across seeds. (D–F) Lowest energy gap in the final batch of separate runs without trace recording. Each point represents one seed.

To test how solver choice affected phasing, we compared CFC and LQA across all 22 autosomes using 100 candidates, 3,000 iterations and five seeds. Mean HE was 0.534% for CFC and 0.594% for LQA; both gave a mean SE of 0.259%. Candidate generation took 0.65 s per chromosome with CFC and 0.61 s with LQA.

### 3.5. Quantum circuit simulation

We compared ideal single-layer QAOA with uniform random sampling on 44 chr22 components with at most 21 variables each. With 512 samples per component, QAOA found the exact minimum energy in 42.6 of the 44 components on average, compared with 34.3 for uniform sampling (Table 2). QAOA also had a smaller total energy gap: 8.287 versus 157.645.

**Table 2.** Recovery of exact minimum energies on 44 chr22 components.

| Sampler | Runs | Minima found / 44 | Total gap |
|---|---|---|---|
| Ideal QAOA | 20 | 42.6 | 8.287 |
| Uniform | 20 | 34.3 | 157.645 |
| Kingston pilot | 1 | 42 | 29.920 |

Each sampler returned 512 samples per component. For each component, we subtracted the exact minimum energy from the lowest sampled energy and summed these gaps across all 44 components. Simulation results are means over 20 seeds; the Kingston pilot was one hardware run.

### 3.6. Chromosome-scale execution on quantum hardware

We constructed a long-read graph for each of the 22 HG002 autosomes and ran the compressed graphs on quantum hardware one connected component at a time. Across the 22 graphs, 3,904 components contained at least one edge. These components represented 1,732,543 of the 2,179,974 observed SNPs (79.5%); the remaining SNPs belonged to isolated phaselets (Figure 5A).

**Figure 5.**
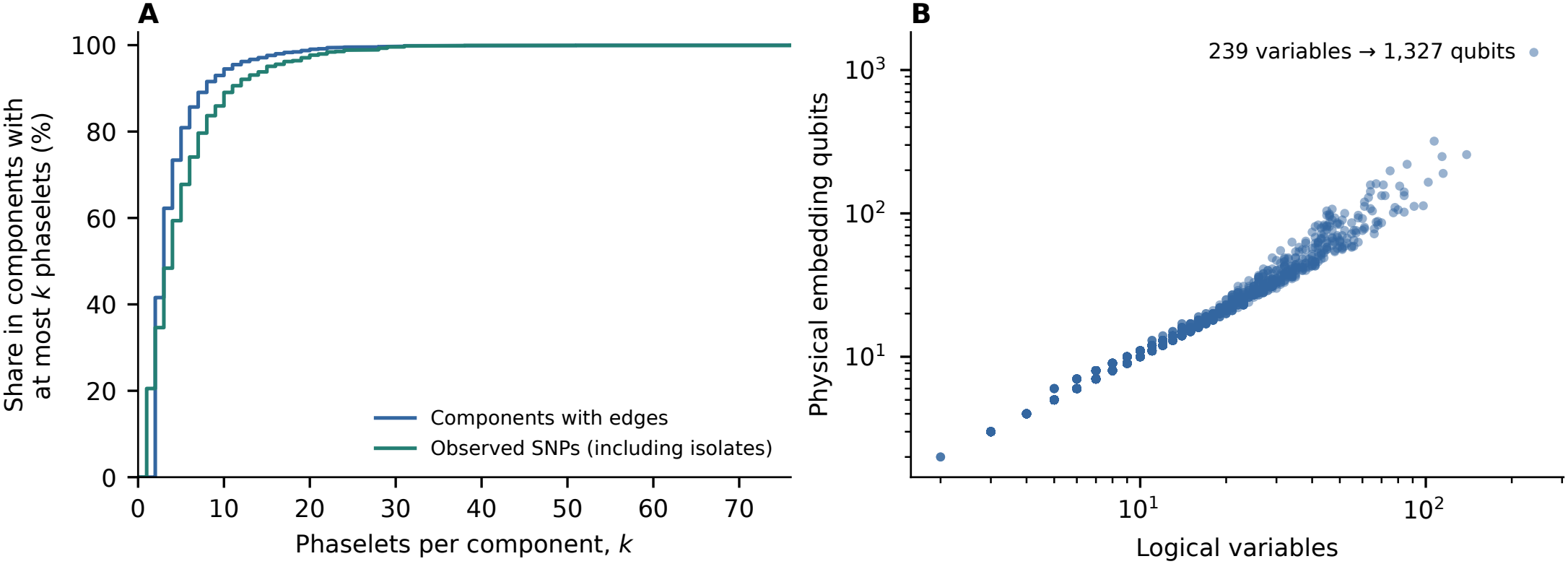
Component sizes and D-Wave qubit use across the 22 HG002 autosomes. (A) The blue curve shows the fraction of components with at most *k* phaselets, counting only components with edges. The green curve shows the fraction of observed SNPs in components with at most *k* phaselets, including SNPs in isolated phaselets. (B) Each point shows the number of logical variables in one component and the number of physical qubits used to embed it on D-Wave.

The D-Wave annealer (`Advantage2_system1`) sampled all 3,904 components. The largest component had 239 logical variables and used 1,327 physical qubits after embedding (Figure 5B). On IBM

Fez, single-layer QAOA sampled 3,903 components with at most 139 problem qubits per circuit. The only component omitted on IBM was the 239-variable component on chr19, leaving its 253 observed SNPs unphased. Both platforms returned 512 samples per component. D-Wave reported 377.35 s of QPU access time, and IBM reported 581 s of charged QPU time for the 22 autosomal jobs.

Across all 22 autosomes, mean HE was 0.584% for D-Wave and 0.548% for CFC on the same graphs; mean SE was 0.259% and 0.258%, respectively. To include IBM in the comparison, we excluded the 253 chr19 SNPs and evaluated all three outputs on the same sites and phase sets (Table 3). On these sites, D-Wave and CFC gave similar error rates, while IBM QAOA had higher mean HE (3.419%) and SE (0.304%).

**Table 3.** Phasing errors on the same sites and phase sets across the 22 HG002 autosomes.

| Sampler | HE (%) | SE (%) |
|---|---|---|
| GPU (CFC) | 0.544 | 0.252 |
| D-Wave annealing | 0.580 | 0.254 |
| IBM QAOA | 3.419 | 0.304 |

Values are means across chromosomes. For each component, we selected the lowest-energy assignment from 100 CFC candidates or 512 hardware samples. D-Wave and CFC predictions were unchanged; the chr19 sites were excluded only from evaluation.

To measure variation between hardware runs, we sampled the same 51 chr22 components three times on each of three IBM processors: Fez, Marrakesh and Kingston. All nine jobs used the same QAOA angles and 512 shots per component. HE ranged from 1.467% to 5.403% across jobs (mean 3.386%), while SE ranged from 0.299% to 0.348% (mean 0.316%).

## 4. Discussion

Many long reads support the same local phase relationships. Phirefly fixes the relative phases of well-supported SNP pairs and uses reads that span the resulting groups to orient them. This allows us to run phasing graphs from all 22 HG002 autosomes on quantum hardware, one connected component at a time.

Choosing which SNP pairs to fix is a heuristic. If a pair is fixed in the wrong relative phase, flipping the whole group cannot correct it. Phirefly releases hard links at sites where the other SNPs on the same reads favor the opposite phase. Across the 22 autosomes, release reduced mean Hamming error but slightly increased mean switch error. Grouping reads is also an approximation: reads that agree on relative phase can differ in the strength of their support, so combining them can change which assignment scores best.

Compression made phasing problems from all 22 HG002 autosomes small enough to run on quantum hardware. On the same sites and phase sets, D-Wave gave phasing error rates close to those of the GPU solver, while single-layer QAOA on IBM gave higher error rates. The same compression also makes many components easy to solve classically. With larger processors, we could fix fewer SNP pairs before sampling and compare quantum and classical solvers on the resulting graphs. This would test whether leaving more phase relationships to the solver improves phasing accuracy and at what computational cost.

## Supporting information

Supplementary methods and results

## Availability and implementation

Source code, installation instructions, example datasets and reproducible benchmark commands are available at https://github.com/wangyf1125/Phirefly.

## Data availability

We used GIAB HG002 PacBio HiFi Revio 20231031 reads and the GRCh38 v5.0q draft variants. The supplementary records list the sources and checksums for the input files, including all 16 Hi-C FASTQ files.

## Author contributions

Yifei Wang developed the methodology, implemented the software and drafted the manuscript. Heng Li, Megan K Le contributed to methodological development, drafted the manuscript and supervised the work.

## Acknowledgements

We thank Haoyu Cheng for his guidance and helpful discussions on haplotype phasing and quantum computing. Y.W. also thanks him for his mentorship and support throughout this work. We also thank Daniel Blankenberg and Fabio Cumbo for helpful discussions on quantum computing, and Fabio Cumbo for testing Phirefly and evaluating QAOA performance through simulations.

