## Supplementary methods and results for "Phirefly: hierarchical quantum haplotype phasing at the chromosome scale"

### S1 Evaluation

We matched SNPs by chromosome, position and alleles and measured Hamming error (HE) and switch error (SE) with WhatsHap 2.8. Input completeness is the fraction of input heterozygous SNPs reported as phased, including SNPs in single-site phase sets.

For comparisons on the same sites and blocks, we kept only input SNPs phased by every method and the reference. For HE, we grouped SNPs together only if every output and the reference placed them in the same block. We assessed blocks with at least two SNPs. For SE, we used only pairs that remained adjacent within an evaluation block for every method after restricting to these sites.

For phase-set span N50, we measured each phase set with at least two SNPs from its first to its last SNP, including both endpoints. For PB-N50 and PB-NG50, WhatsHap splits overlapping blocks and measures each block as the last position minus the first. PB-N50 uses half the total block length as its threshold; PB-NG50 uses half the chromosome length and is zero if the blocks do not reach it. We gave each chromosome equal weight when reporting means.

### S2 Link filtering and compression

Each read contributed one vote for the relative phase of a SNP pair. For SNPs $i$ and $j$, let $n_{ij}$ be the number of contributing reads, $K_{ij}$ the sum of their signed votes and $d_{ij}$ the distance in base pairs. With unit weights, the filtering score is

$$\rho_{ij} = 0.55\left(1 - \frac{|K_{ij}|}{n_{ij}}\right) + \frac{0.15}{\ln(1+n_{ij})} + 0.10\min(d_{ij}/10^6, 1) + \frac{0.20}{n_{ij}}. \tag{S1}$$

We applied the hard-link filter first and tested the remaining links against the soft-link filter (Table S1). Releasing a hard link leaves its read observations in the model but does not turn it into a soft link (Figure S1). We capped each soft link before summing links between the same two phaselets, so the sum can exceed 10.

Combining reads can change their score even before rescaling. For example, at phaselet orientations $(+1, -1)$, two reads with projected weights $(2, 1)$ and $(1, 2)$ contribute $|2-1| + |1-2| = 2$ separately. Their combined weights are $(3, 3)$, which give $|3-3| = 0$.

Table S1: Default settings for long-read phasing.

| Setting | Hard links | Soft links |
|---|---|---|
| Minimum contributing reads | 2 | 1 |
| Maximum distance | 1 Mb | 1 Mb |
| Maximum filtering score $\rho_{ij}$ | 0.45 | 0.75 |
| Additional requirement | $K_{ij}\tau_i\tau_j > 0$ | $\|K_{ij}\|/n_{ij} \geq 0.55$ |

Minimum base quality 13; minimum mapping quality 20; unit observation weights.
Soft links: multiply by 0.25 and cap the magnitude at 10 before summation.
Hyperreads: divide by the square root of the group size and cap the magnitude at 10.
CFC: 100 candidates, 1,000 iterations, time step 0.1, seed 23.
CFC initial scale 0.001; Gaussian perturbation 0.003; single-precision arithmetic.

### S3 Runtime measurement

We ran each method three times per chromosome. Each run started a new process, and we randomized method order within each chromosome and repetition. HapCUT2 used a minimum base quality of 4 in PacBio mode; Phirefly and QHap SNP used 13. Phirefly and QHap SNP used eight workers for read extraction and one thread for numerical libraries. We turned off debug output, convergence tracing and memory polling during timing.

For WhatsHap, Core time covered dynamic programming and traceback; for HapCUT2, it covered random initialization and Max-Cut optimization. For QHap SNP, we timed all ballistic simulated bifurcation calls; for Phirefly, we timed CFC. GPU solver time included initialization, data transfers, updates and scoring within the solver, sample readback and synchronization. Read extraction, graph construction, final sample selection and VCF export counted only toward end-to-end time.

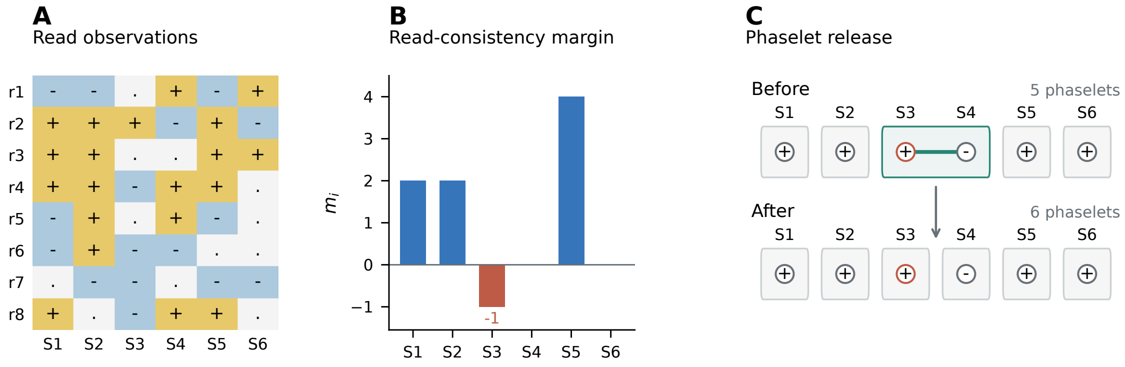


Figure S1: Releasing a hard link. (A) Alleles observed on each read; dots mark missing observations. (B) To check a SNP, we assign each read using its other SNPs and calculate the margin at the site being checked. (C) S3 has a negative margin. Removing the S3–S4 link lets the solver orient the two SNPs independently.

#### S4 Phasing controls

Changing only the CFC seed gave mean HE of 0.534–0.603% and mean SE of 0.257–0.258% (Table S2). To test release, we paired runs with and without release by chromosome and seed. Release lowered mean HE from 1.000% to 0.551% and raised mean SE from 0.253% to 0.258% (Table S3).

Table S2: Effect of the CFC seed on phasing error. Each run used 100 candidates and 1,000 iterations. Values are means across the 22 HG002 autosomes.

| Seed | 1 | 2 | 3 | 4 | 23 |
|---|---|---|---|---|---|
| Mean HE (%) | 0.534 | 0.570 | 0.555 | 0.603 | 0.548 |
| Mean SE (%) | 0.257 | 0.258 | 0.258 | 0.258 | 0.258 |

Table S3: Phasing with and without release. Each comparison used the same SNPs and evaluation blocks for HE and the same adjacent pairs for SE. Values are means over 22 chromosomes and three seeds (1, 2 and 23).

| Setting | HE (%) | SE (%) |
|---|---|---|
| Without release | 1.000 | 0.253 |
| With release | 0.551 | 0.258 |

Table S4 compares four models on the same SNPs in each of three regions. The raw model uses one variable per read and one per SNP. The phaselet model projects reads onto SNP groups. The grouped model also combines reads that link the same groups with the same relative signs. The full model adds group-size scaling, clipping and soft links together. Compared with grouping alone, these changes lowered HE in the chr20 interval but raised it in the MHC and chr19 intervals.

Table S4: Effect of compression on model size and phasing error. Each model used 100 candidates, 1,000 updates and seeds 1, 2 and 23. Within each interval, all four models used the same observed SNPs and evaluation blocks. HE and SE are percentages averaged over seeds. GRCh38 intervals: MHC, chr6:28,510,120–33,480,577; chr20:10,000,000–11,000,000; chr19:8,595,901–8,775,840.

| | MHC | | | chr20 interval | | | chr19 interval | | |
|---|---|---|---|---|---|---|---|---|---|
| Model | Variables | HE | SE | Variables | HE | SE | Variables | HE | SE |
| Raw | 22,459 | 1.392 | 0.246 | 3,910 | 7.669 | 0.419 | 817 | 46.377 | 26.323 |
| Phaselet | 1,209 | 0.176 | 0.236 | 97 | 0.342 | 0.686 | 331 | 35.968 | 27.381 |
| Grouped | 187 | 0.176 | 0.236 | 11 | 0.342 | 0.686 | 239 | 20.553 | 26.190 |
| Full | 187 | 0.199 | 0.269 | 11 | 0.228 | 0.457 | 239 | 35.968 | 25.000 |

For the seven-solver comparison, we found the exact minimum of the compressed chr22 model by enumerating phaselet orientations and choosing the best label for each hyperread at each assignment. For chr1 and chr20, the reference was the lowest energy found by any solver.

### S5 Quantum execution

For each component, let $J_{\max} = \max_{u<v} |J_{uv}|$ be its largest absolute coupling and $n$ its number of variables. The single-layer QAOA circuit prepares

$$|\psi(\beta,\gamma)\rangle = e^{-i\beta \sum_u X_u} e^{-i\gamma \widehat{H}_C / J_{\max}} |+\rangle^{\otimes n}, \qquad \widehat{H}_C = -\sum_{u<v} J_{uv} Z_u Z_v. \tag{S2}$$

Here $X_u$ and $Z_u$ are Pauli operators on qubit $u$; reported energies use the original coupling scale. We minimized the analytic expected energy with SciPy differential evolution over $\beta \in [-\pi/4, \pi/4]$ and $\gamma \in [0, \pi]$, using seed 23, a population multiplier of 10, at most 80 generations, tolerance $10^{-8}$ and final polishing with one worker. We compiled circuits at optimization level 3 with transpiler seed 23. Measurement bits 0 and 1 map to spins +1 and −1, respectively. Ideal simulation used Aer statevectors; the uniform control assigned each spin +1 or −1 independently with equal probability.

On IBM, ties for the lowest sampled energy were resolved by storage order on chr1–21 and lexicographic bitstring order in the chr22 repetitions. The autosomal comparison used the first Fez run for chr22; the Kingston pilot was separate from the nine repeated jobs. For the three-sampler comparison, we excluded the same 253 observed SNPs from chr19:8,595,901–8,775,840 without changing the D-Wave or CFC assignments.

On D-Wave, we found embeddings with `minorminer` (seed 23) and set chain strength by uniform torque compensation with prefactor 1.414. We used automatic coefficient scaling and decoded chains by majority vote, without further chain repair. The adapter used `dimod` 0.12.23, `dwave-system` 1.36.0 and `minorminer` 0.2.22.

### S6 Hi-C integration

We aligned 16 HG002 HPRC_PLUS Hi-C FASTQ files with BWA 0.7.19-r1273. We excluded secondary, supplementary, duplicate and QC-failing alignments and required mapping quality at least 20 and base quality at least 13. Bases with missing qualities were excluded. Within each alignment, we chose the phaselet with the largest absolute signed allele sum $|a|$; a zero sum gave no anchor. The anchor sign was $\mathrm{sign}(a)$ and its weight was $q = |a| \max(\min(\mathrm{MQ}, 60)/60, 0.05)$.

For each read name after normalization, we retained the strongest anchor per phaselet. The read name contributed at most once to each phaselet pair, with weight $\max(0.1, \min(q_k, q_\ell))$ and sign equal to the product of the anchor signs. Each retained pair required at least one read name. For links between different long-read components, we also required at least 20 external contact names at each endpoint component. We summed signed weights for each pair, multiplied by 0.1 and capped the result at $[-10, 10]$. For candidate selection, we kept phaselets connected by long-read or Hi-C links in the same component even when their coefficients cancelled. HapCUT2 used `-nf 1` for mixed-technology fragments and `-hic 1` for Hi-C extraction and phasing.

Table S5: Hi-C phasing across the 22 HG002 autosomes. The first three rows use each method's own phase sets. The final two use identical evaluation blocks for HE and identical adjacent pairs for SE (2,177,900 SNPs and 2,173,404 pairs in total). Values are chromosome means; block lengths describe the original outputs.

| Method | HE (%) | SE (%) | PB-N50 (Mb) | Phase-set span N50 (Mb) |
|---|---|---|---|---|
| *Each method's own phase sets* | | | | |
| Phirefly, long reads | 0.548 | 0.258 | 0.564 | 0.564 |
| Phirefly + Hi-C | 0.207 | 0.214 | 3.448 | 125.999 |
| HapCUT2 + Hi-C | 0.267 | 0.104 | 42.755 | 115.052 |
| *Same SNPs and evaluation blocks* | | | | |
| Phirefly + Hi-C | 0.205 | 0.212 | – | – |
| HapCUT2 + Hi-C | 0.120 | 0.099 | – | – |
